# INTEROCEPTIVE INFORMATION OF PHYSICAL VIGOR THROUGH CIRCULATING INSULIN-LIKE GROWTH FACTOR 1

**DOI:** 10.1101/2021.05.25.445442

**Authors:** Jonathan A. Zegarra-Valdivia, M. Zahid Kahn, Jansen Fernandes, Julio Esparza, Kentaro Suda, M. Estrella Fernandez de Sevilla, Sonia Díaz-Pacheco, Ignacio Torres Aleman

## Abstract

The brain relies on interoceptive feedback signals to regulate bodily functions. Mice with low serum IGF-1 levels (LID mice) exhibit reduced spontaneous running, a behavior that normalizes after sustained systemic IGF-1 treatment. This observation led us to hypothesize that circulating IGF-1—a key regulator of skeletal muscle and bone mass that crosses the blood-brain barrier during physical activity—may convey body vigor information to the brain. Since hypothalamic orexin neurons, that are involved in regulating physical activity, express IGF-1 receptors (IGF-1R) and are modulated by this growth factor, we hypothesized that these neurons might gauge circulating IGF-1 levels to modulate physical activity. Indeed, inactivation of IGF-1R in mouse orexin neurons (Firoc mice) was associated to less time spent in free running. These mice maintain physical fitness but display altered mood and are less sensitive to the rewarding actions of exercise. Further, in response to exercise, Firoc mice showed limited c-fos activation of hypothalamic orexin neurons and monoaminergic neurons of the ventro-tegmental area (VTA) in the brainstem. This area is involved in the rewarding component of exercise that seems to be modulated by IGF-1, as mice receiving systemic IGF-1 showed increased c-fos expression in VTA neurons, while mice with reduced IGF-1R expression in VTA neurons showed no improved mood after exercise. Collectively, these results suggest that circulating IGF-1 is gauged by orexin neurons to modulate physical activity, and that VTA neurons convey the rewarding properties of exercise through direct actions of IGF-1 on them. Hence, serum IGF-1 may constitute an interoceptive signal acting onto orexin/VTA neurons to modulate physical activity according to physical vigor (muscle and bone mass).

## Introduction

Information about the body’s internal state is transmitted to the brain via interoceptive pathways, comprising humoral factors and the peripheral nervous system (Craig, 2002). Many body-to-brain signals, that are likely involved in modulation of brain states (Flavell et al., 2022), have been extensively analyzed concerning their cellular targets, actions, and circuits involved (Critchley et al., 2004; Allen, 2020). These include those affecting hunger/satiety, adiposity, posture/muscle tone, visceral function and others (Quigley et al., 2021). Interoceptive signals conveying muscle strength (vigor) to brain centers are considered the sum of visceral, musculoskeletal, and sensory inputs, but they may also include humoral factors such as insulin-like growth factor 1 (IGF-1). Indeed, IGF-1 has been proposed to sustain apparent vigor, based on its numerous actions in health and disease (Ayres, 2020). Circulating IGF-1, mainly derived from the liver (Yakar et al., 1999), is taken up by the brain in response to exercise (Carro et al., 2000), and displays a broad range of effects potentially related to vigor awareness, such as modulation of mood (Santi et al., 2018), which contributes to wellbeing states. Moreover, IGF-1 is secreted into the circulation in response to muscle activity (DeVol et al., 1990; Berg et al., 2007), is vital in maintaining muscle (Clemmons, 2009) and bone mass (Yakar et al., 2002; Wang et al., 2023), and reflects muscle strength (Bucci et al., 2013). Thus, circulating IGF-1 is a potential candidate to inform of muscle/bone status to brain centers in charge of controlling physical activity, a main component of behavioral expression of internal states (Flavell et al., 2022).

In this vein, we hypothesized that hypothalamic orexin neurons sense the entrance of serum IGF-1 following increased physical activity (Carro et al., 2000). Significantly, orexin neurons express IGF-1 receptors, and their activity is regulated by this growth factor (Zegarra-Valdivia et al., 2020; Fernández de Sevilla et al., 2022). These neurons are involved in multiple basic homeostatic mechanisms, coupling for instance waking with muscle tone (Burgess and Peever, 2013), and controlling general physical activity (Kiwaki et al., 2004; Kotz, 2006). Further, orexin neurons are involved in motivational behaviors (Mahler et al., 2014), and physical activity acts as a rewarding stimulus (Greenwood et al., 2011).

Based on the above considerations, we postulate that IGF-1 input to orexin neurons conveys physical vigor information. In turn, orexin neurons would sustain physical activity by modulating muscle tone and conveying vigor status to reward areas such as the brainstem ventral tegmental area (VTA) (Medrano et al., 2021), to positively modulate physical activity. In this way, physical fitness would be associated to wellbeing. We now report that IGF-1 is gauged by orexin neurons, which in turn couple motivation with physical activity by signaling onto the VTA, which is also directly targeted by IGF-1.

## Results

### Serum IGF-1 modulates physical activity

Mice with low serum IGF-1 levels (LID mice, Figure 1A) display a disparity of brain deficits and abnormal behaviors, as characterized in detail before (Trejo et al., 2007; Zegarra-Valdivia et al., 2019). We assessed spontaneous activity in LID female mice by housing them in cages with free access to a running wheel and found that they spent less time running, as compared to control littermates (Figure 1B). We use only females in these series of experiments because wild type female mice displayed significantly higher spontaneous running activity than males (Suppl Fig. 1A, B). This agrees with previously reported sex differences in physical activity (Bartling et al., 2017), and allowed us to better assess spontaneous running. LID mice have been extensively characterized before (Yakar et al., 1999; Sjogren et al., 2002), and they show normal activity levels when placed in an activity cage (Trejo et al., 2007). Since chronic treatment with a subcutaneous infusion of IGF-1 using osmotic mini-pumps normalized almost completely the levels of spontaneous running (Figure 1C), it seems that systemic IGF-1 is involved in regulation of physical activity.

**Figure 1:**
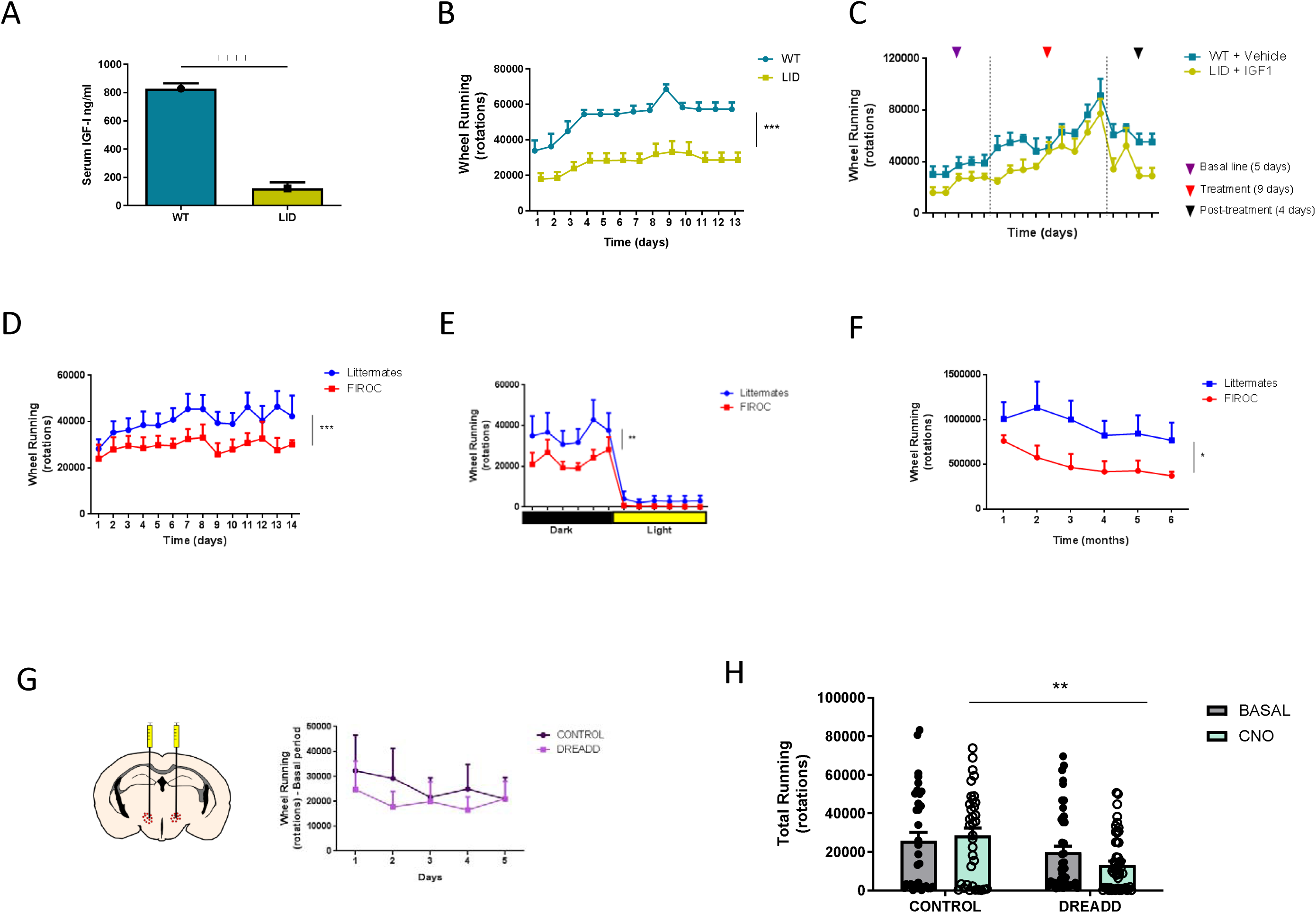
Serum IGF-1 regulates physical activity through orexin neurons. **A,** LID mice have reduced serum IGF-1 levels compared to controls (wild type; WT=6, LID=12; ***p<0.0001, Unpaired t-test, sex balanced) **B,** Female LID mice show reduced spontaneous activity when housed with free access to a running wheel (WT=5, LID=10; ***p<0.0001, Two-Way ANOVA). **C,** Differences in running behavior between WT and LID disappear after one-month chronic treatment with IGF-1 using osmotic mini-pumps (WT=4, LID=4; p=0.1131, Two-Way RM ANOVA). **D,** Female mice lacking functional IGF-1R in orexin neurons (Firoc mice) show reduced spontaneous activity compared to control littermates (WT=10, Firoc=9; ***p<0.0001, Two-Way ANOVA). **E,** Firoc mice show reduced activity only during the active phase (WT=8, Firoc=7; **p=0.0024, Two-Way ANOVA). **F,** Firoc mice show reduced activity for at least 6 consecutive months (WT=5, Firoc=5; *p=0.0358, Two-Way RM ANOVA). **G,** Comparison between DREADDi-Orexin-Cre mice and controls (mCherry virus) did not show differences in running under basal conditions (without CNO). Left: representative diagram of coronal section indicating the LH area where double-injection of DREADDi-expressing viruses was carried out. **H,** Chemogenetic inhibition with CNO of orexin neurons in Orexin-Cre mice (n=50) leads to reduced spontaneous activity, as measured in the running wheel for 1 hour of free access to it. Control mice received a control mCherry virus. CNO was given 45 min before the running wheel was made accessible to the mice (n=43; **p=0.0018, Two-Way ANOVA).

Next, we analyzed orexin neurons as possible targets of circulating IGF-1 since these neurons are involved in regulation of physical activity (Kosse et al., 2017), and we already showed that they are modulated by IGF-1 (Zegarra-Valdivia et al., 2020; Fernández de Sevilla et al., 2022; Pignatelli et al., 2022). We observed that female mice lacking IGF-1 receptors in orexin neurons (Firoc mice) also showed diminished spontaneous activity (Figure 1D). Firoc mice show increased sleepiness during the inactive, but not the active phase of the day, ruling out a secondary consequence of altered sleep architecture previously characterized by us in these mice (Zegarra-Valdivia et al., 2020). Indeed, decreased activity is concentrated in the active phase in Firoc mice (Figure 1E), as also seen in LID mice (Suppl Fig. 1C). Reduced spontaneous activity in Firoc mice was consistent along time, as they showed this deficit for at least 6 months (Figure 1F).

Since activation of orexin neurons has been shown to be required for spontaneous physical activity (Kosse et al., 2017), we inhibited them by bilateral injection of DREADDi-expressing Cre-dependent viruses in the lateral hypothalamus (AP= -1.4; ML= ± 0.9; DV= -5.4) of Orexin-Cre mice (Figure 1G). No differences in basal running activity between control mice receiving a mCherry-expressing control virus and DREADDi-injected mice were found during the five days prior to CNO administration (Figure 1G). On the test day, CNO was given 45 min before opening for 1 hour the access to the running wheel. While viral DREADD injection *per se* reduced total running time (basal), only CNO administration significantly reduced spontaneous physical activity in DREADDi-Orexin-Cre mice, as compared to DREADDi-injected controls receiving the vehicle (Figure 1H).

Using c-Fos immunostaining as a marker of neuronal activity (Zhang et al., 2002), we confirmed that exercise activates orexin neurons (Suppl. Fig. 1D-F), consistent with prior findings (Martins et al., 2004). We next examined whether this activation also occurs in Firoc and LID mice (Figure 2). When Firoc mice were given 1-hour free access to a running wheel, we observed a significant reduction in the number of orexin^+^/c-Fos^+^ double-labeled cells in the lateral hypothalamus (LH) 2 hours post-exercise (Fig. 2A,B,E). This suggests that circulating IGF-1 may play a role in exercise-induced orexin neuron activation, a hypothesis further supported by results in LID mice subjected to the same running protocol. Like Firoc mice, LID mice exhibited fewer orexin^+^/c-Fos^+^ cells in the LH (Fig. 2C,D,F). Importantly, the total number of orexin neurons did not differ significantly between Firoc, LID, and their respective controls (Suppl. Fig. 2A,B).

**Figure 2:**
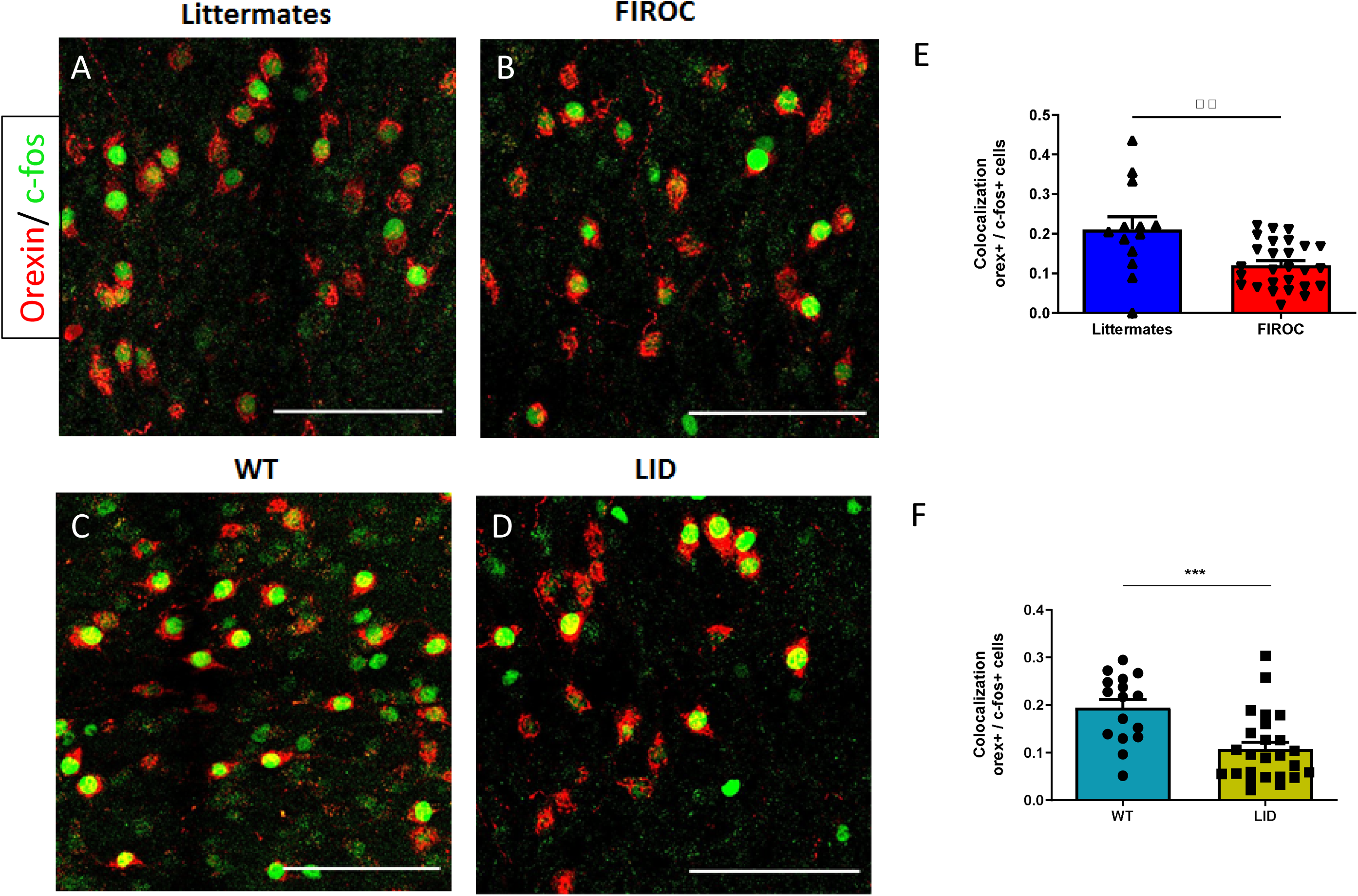
Running elicits reduced activation of orexin neurons in LID and Firoc mice. **A-C,** Both Firoc (A,B) and LID (C,D) mice submitted to voluntary wheel running show a reduced number of double-labeled orexin (red)/c-fos (green) cells in the lateral hypothalamus as compared to controls (kept in running cages with locked wheels). Bars are 50 µm. **E,** Quantification histograms of double-labeled orexin/c-fos neurons in littermate controls (n=27 fields area of LH, 3 mice) and Firoc mice (n=13 fields area of LH, 3 mice) show significant differences (*p=0.0167, Unpaired t-test, Welch correction). **F,** Quantification histograms of double-labeled orexin/c-fos neurons in WT (n=16 fields area of LH, 3 mice) and LID (n=25 field area of LH, 3 mice) show significant differences (***p<0.001, Unpaired t-test).

### IGF-1 modulates physical activity through orexin and VTA neurons

Since Firoc mice show unaltered body weight (Suppl Fig 2C) and muscle mass (Suppl Fig 2D,E), normal strength (Suppl Fig 2F,G), intact motor coordination (Suppl Fig 2H), and similar physical activity than control littermates (Suppl Fig 2I), we analyzed their mood status to determine whether they present motivational disturbances that could explain their lower level of physical activity, as orexin is involved in motivated behaviors (Sakurai, 2014). Behavioral analysis of Firoc mice (see behavioral testing procedure in Suppl Fig 3A) evidenced reduced resilience to stress and increased anxiety-like behavior, as determined by longer immobility time in the forced swim test and shorter time visiting the open arms of the elevated plus maze, respectively (Figure 3A, B). Interestingly, anxiety levels were normal in younger Firoc mice (not shown), suggesting that mood disturbances develop gradually in these mice. In addition, Firoc mice presented normal social affiliation but a reduced social novelty/preference behavior (Figure 3C, D). The latter indicates reduced motivation and suggests reduced rewarding effects of social interactions. In LID mice resilience to stress was normal (Figure 3E), while anxiety levels were increased (Figure 3F). LID mice also present normal social affiliation and reduced social novelty/preference (Figure 3G, H). Thus, the behavioral pattern of LID mice was similar, but not identical to Firoc mice.

**Figure 3:**
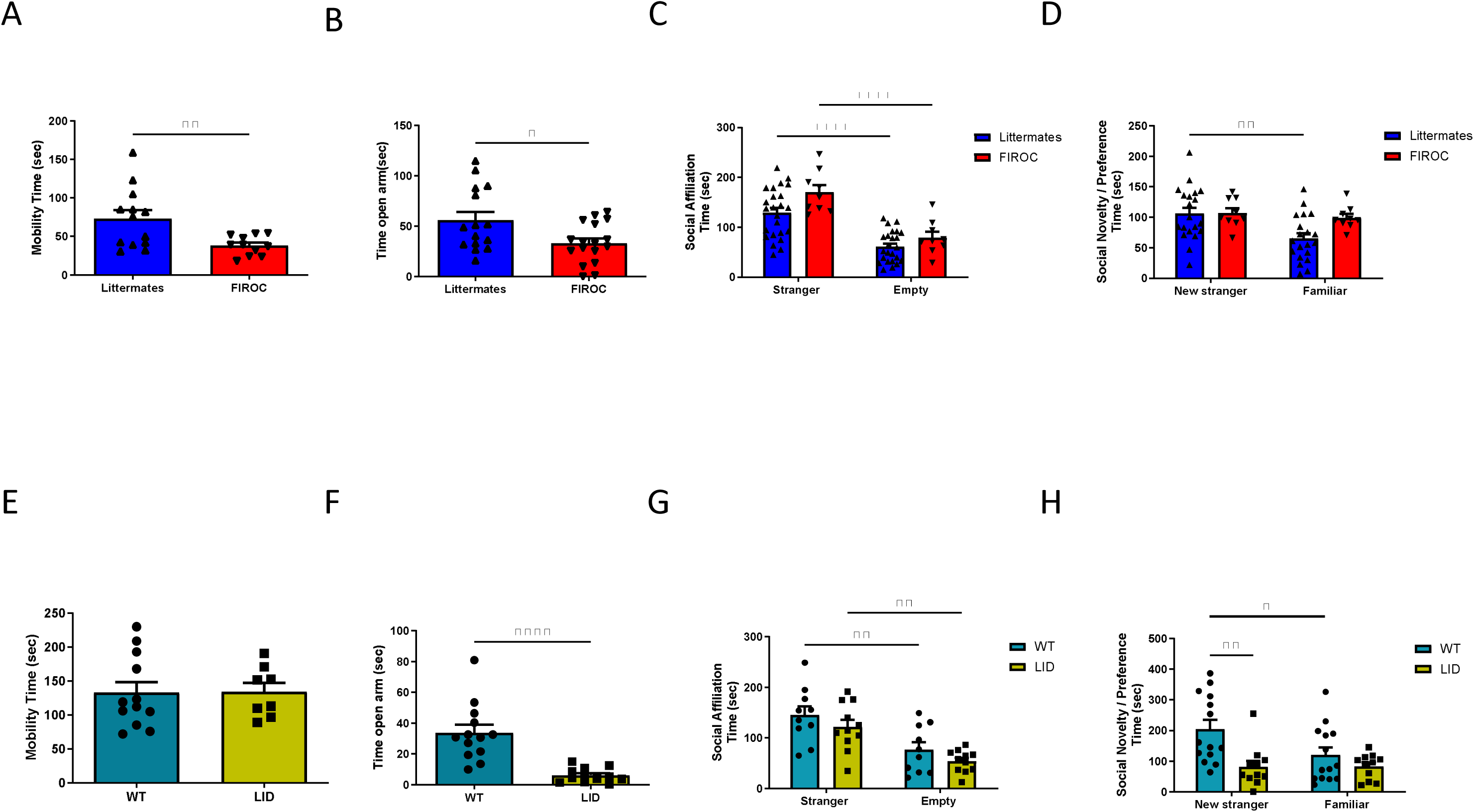
Behavioral disturbances in Firoc and LID mice. **A,** Depressive-like behavior in Firoc mice, as determined in the forced swim test, was increased (WT=11, Firoc=13; **p<0.001, Unpaired t-test). **B,** Increased anxiety in Firoc mice, as determined in the elevated plus-maze (WT=17, Firoc=15; *p=0.0177, Unpaired t-test). **C,** Social affiliation in Firoc mice was normal, as indicated by the “stranger vs. empty box” test (WT=24, Firoc= 9; ***p<0.001, Two-Way ANOVA). **D**, Social novelty/preference in Firoc mice was disrupted, as indicated by the “new-partner vs. familiar partner” test (WT= 21, Firoc= 9, WT preference **p=0.0085, Firoc preference p=0.9698, Two-Way ANOVA). **E,** As determined in the forced swim test, depressive-like behavior in LID mice was similar in both groups (WT=12, LID=8; p=0.9520, Unpaired t-test). **F,** Increased anxiety is found of LID mice as determined in the elevated plus-maze (WT=13, LID=12; ***p<0.0001, Unpaired t-test, Welch Correction). **F,** Social affiliation in LID mice was normal, as indicated by the “stranger vs. empty box” test (WT=10, LID=11; **p<0.01, Two-Way ANOVA). **G,** Social intercourse is significantly altered in LID mice, as indicated by the “new-partner vs. familiar” test (WT=14, LID=11; WT preference *p<0.05, LID preference p=0.9531, WT vs. LID new stranger preference, **p=0.0042, Two-Way ANOVA).

We next analyzed whether the ventral tegmental area (VTA), that forms part of the reward circuitry (Morales and Margolis, 2017), and is involved in the rewarding aspects of exercise (Medrano et al., 2021) is also involved in the modulatory actions of IGF-1. First, we confirmed previous observations that IGF-1 modulates the activity of dopamine neurons in the VTA (Pristera et al., 2019). After systemic administration of IGF-1 (1 μg/g, ip) to wild-type (WT) mice, we determined 2 hours later the number of monoaminergic (TH^+^) cells expressing c-fos in the VTA (Herrera et al., 2016). IGF-1 significantly increased the number of double-labeled TH^+^/c-fos^+^ cells compared to saline-injected mice (Figure 4A, B), suggesting that it activates these neurons. As seen in orexin neurons, exercise (1 hour running in the wheel, 2 hours before sacrifice) also induced c-fos activity in TH^+^ VTA neurons of WT mice (Suppl Fig. 3B-D). We then analyzed whether Firoc and LID mice show also disturbed activation of TH^+^ VTA neurons in response to exercise as seen in orexin neurons. Indeed, exercised Firoc mice did not increase c-fos^+^ immunostaining in TH^+^ cells to the level seen in littermates (Figure 4C, D). Conversely, exercised LID mice showed a higher number of double labelled c-fos^+^/TH^+^ cells in the VTA compared to controls (Figure 4E, F). When TH^+^ cells were scored, no differences between groups was observed (Suppl Fig 4A-C).

**Figure 4:**
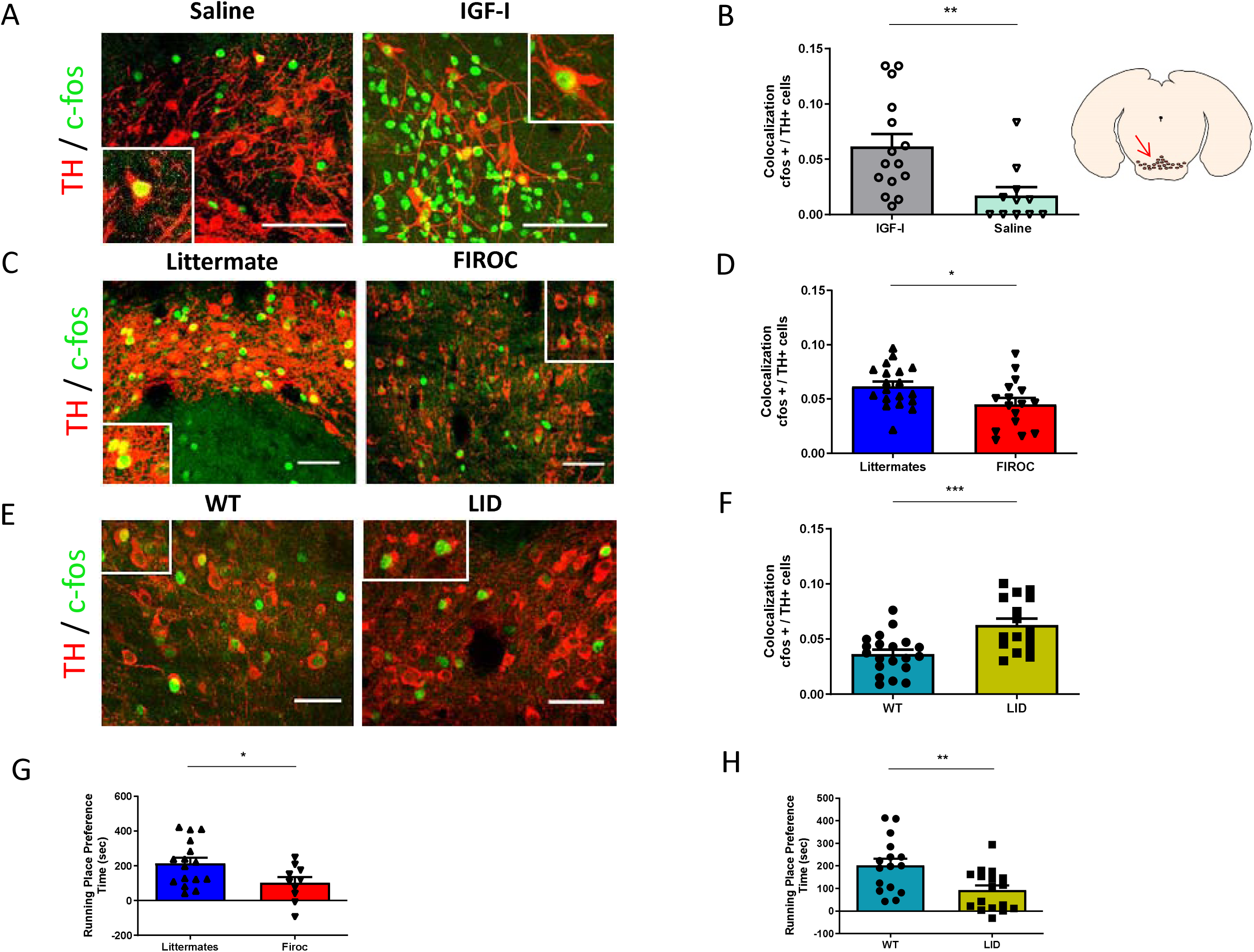
IGF-1 and exercise modulate the activity of monoaminergic VTA neurons. **A,** Representative micrographs of double-stained c-fos (green)/TH (red) cells in the VTA of wild-type mice injected with saline or IGF-1 (1µg/g body weight, ip; two hours before). Bars are 50 µm. **B,** Quantification histograms of double-labelled c-fos^+/^ TH^+^ cells in VTA (saline n=11 fields, 3 mice; IGF-1 mice=15 fields, 3 mice; **p=0.0063, Unpaired t-test). Schematic representation of VTA area (right hand). **C,** Double-stained c-fos^+/^TH^+^ cells in the VTA of Firoc and WT (littermates) after 1 h exercise. Bars are 50 µm. **D,** Quantification histograms of double-labelled c-fos^+^/TH^+^ cells (WT=19 fields, Firoc=16 fields, 3 mice per group; *p=0.0256, Unpaired t-test). **E,** Double-stained c-fos^+^/TH^+^cells in the VTA of LID mice after 1 h exercise. Bars are 50 µm. **F,** Quantification histograms of double-labelled c-fos^+^/ TH+ cells (WT=4 fields, LID=6 fields, 3 mice per group; p=0.1388, Mann-Whitney t-test). **G,** The rewarding effects of exercise are attenuated in Firoc mice, as seen by significantly reduced time spent in the exercise chamber (WT=16, Firoc=10; *p=0.029, Unpaired t-test). H, Place preference is also reduced in LID mice (WT=16, LID=17; **p=0.0049, Unpaired t-test).

Although exercise improved mood in wild-type mice, as evidenced by reduced anxiety (increased time spent in the open arms of the elevated plus maze; Suppl. Fig. 3E), its conditioning effects were attenuated in both Firoc and LID mice. This was demonstrated by the conditioned place preference (CPP) test for running, despite the opposing c-fos activation patterns observed in VTA neurons of these two strains. Thus, both Firoc and LID mice spent significantly less time in the running chamber than control littermates (Figure 4G,H).

Given these findings, we administered the dopamine D1 receptor agonist SKF-82958 (DA1; 0.1 mg/kg, i.p.) to LID and Firoc mice, as dopaminergic signaling in the ventral tegmental area (VTA) plays a critical role in reward processing (Kringelbach and Berridge, 2009). DA1 treatment recovered spontaneous running activity in LID mice (Figure 5A, B), while moderately decreased it in Firoc mice (Figure 5C, D). Since the response to the D1 receptor agonist differs between Firoc and LID mice, and IGF-1 exerts stimulatory actions on c-fos expression in VTA neurons, we first confirmed the presence of IGF-1R in TH neurons in the VTA (Suppl Fig 5A), as reported by others (Pristera et al., 2019), supporting the notion that these neurons are directly targeted by IGF-1. Wed then injected IGF-1R^fl/fl^ mice with a Cre-GFP expressing, synapsin-driven virus in the VTA (Suppl Fig 5B) to downregulate IGF-1R activity in VTA neurons (Suppl Fig 5C). Mice with downregulated IGF-1R expression in VTA neurons showed similar place preference after exercise to that found in control virus injected mice (Figure 6A), indicating that the conditioning actions of exercise were preserved in them. However, exercise did not improve their mood, as determined by no reduced anxiety levels in the EPM after exercise, compared to control virus-injected IGF-1R^fl/fl^ mice (Figure 6B). Sedentary mice show similar anxiety levels regardless of type of virus injected, indicating that viral injection *per se* did not affect this behavior (not shown). Collectively, these observations suggest that the hedonic component of exercise was blocked in the absence of normal IGF-1R activity in VTA neurons.

**Figure 5:**
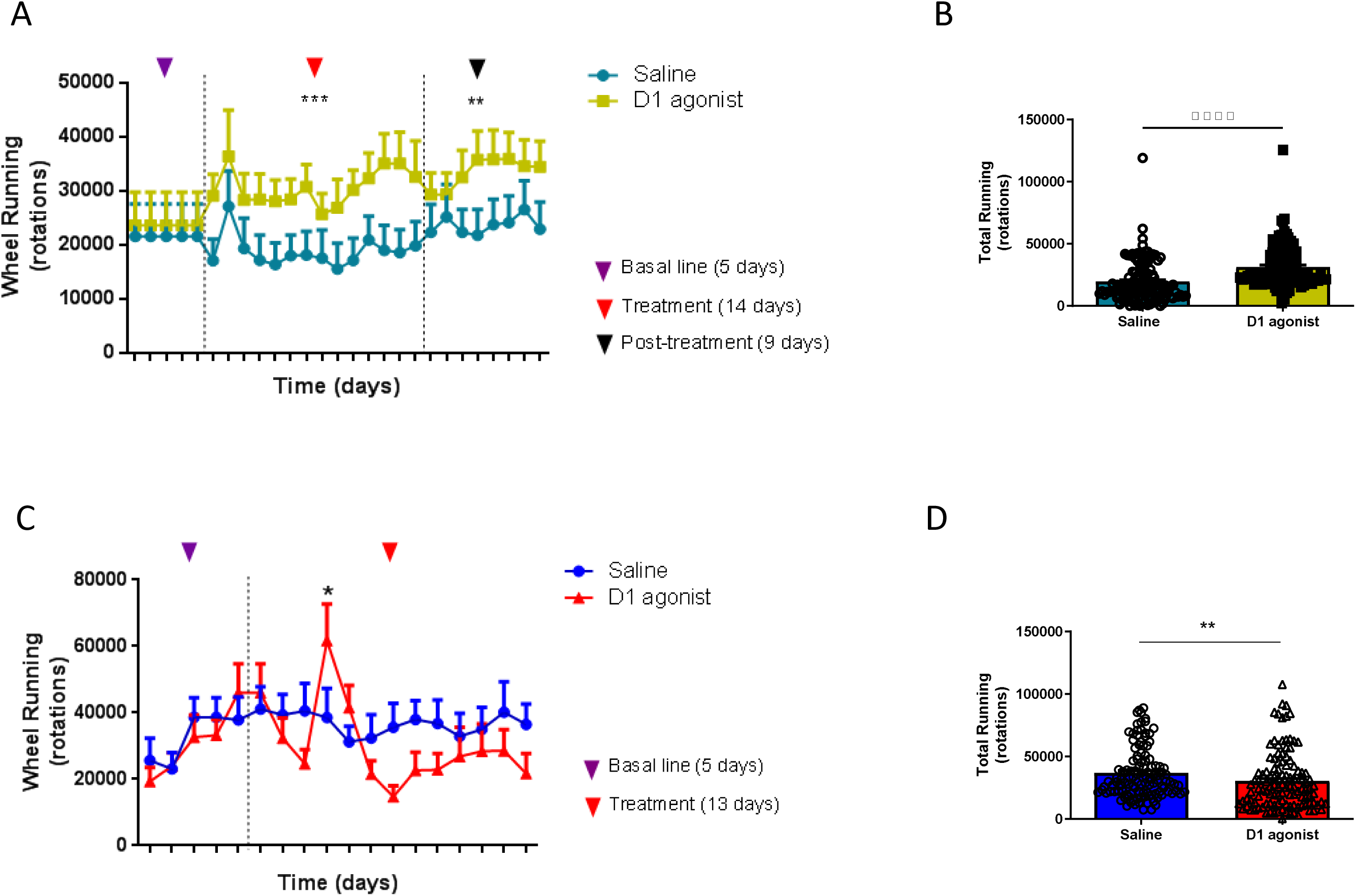
Treatment with a dopamine receptor 1 agonist restores spontaneous physical activity in LID, but not in Firoc mice. **A,** Administration of the D1 agonist SKF82958 (0.1mg/kg, ip) to LID mice significantly increased their spontaneous activity even after treatment ceased (LID saline=10, LID D1 agonist=9; ***p<0.001). **B,** Total activity along time of treatment was increased in SKF82958 -treated LID mice (***p<0.001, Mann-Whitney test). **C,** Firoc mice treated with SKF82958 changed the pattern of spontaneous physical activity (n= 9 per group; p=0.4047, interaction factor=0.0030, day 4 *p=0.0327). **D,** Total activity along time of treatment was decreased in SKF82958 -treated Firoc mice (**p=0.0018).

**Figure 6:**
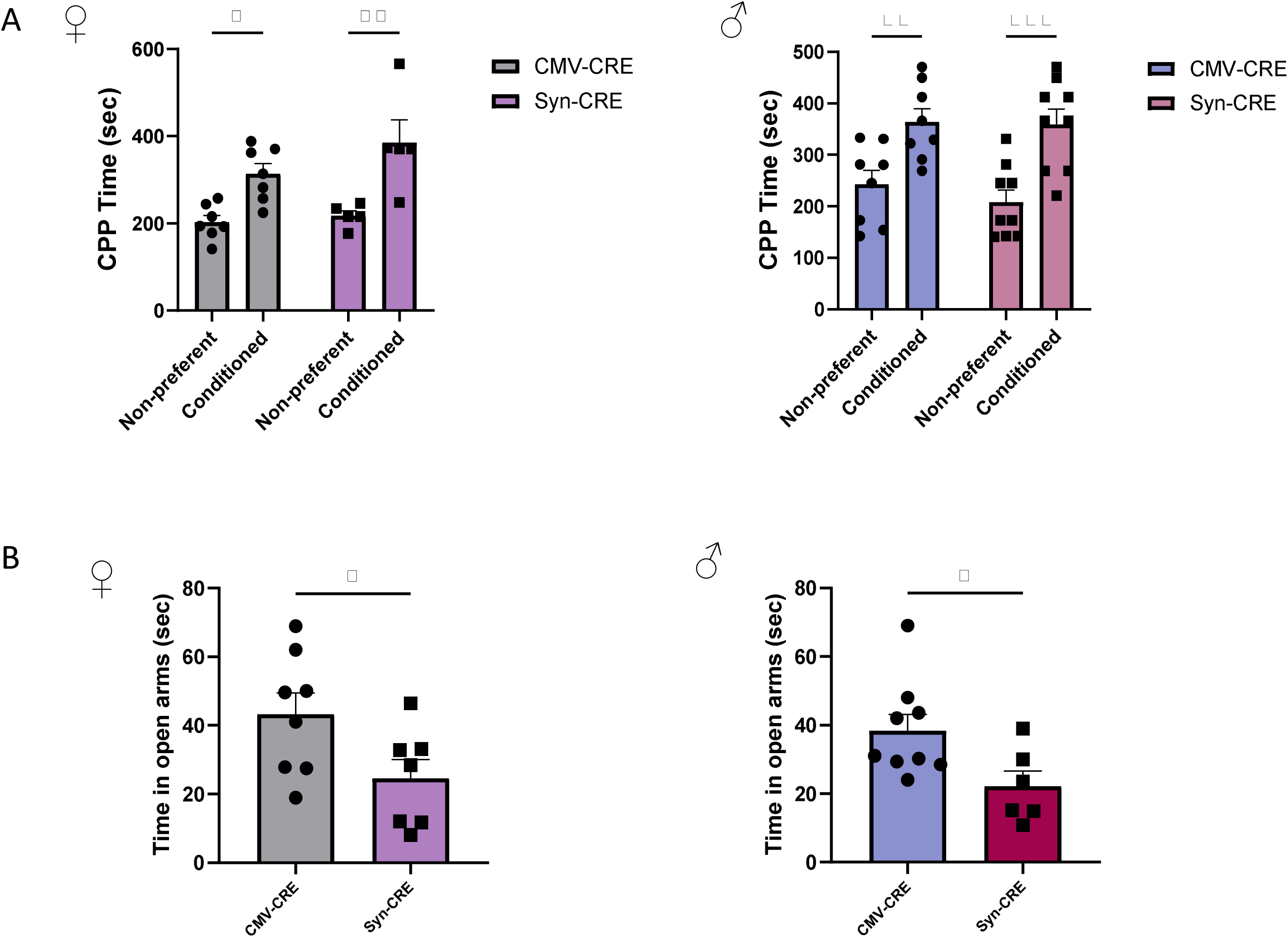
IGF-1 modulates spontaneous activity through the ventral tegmental area. **A,** IGF-1R^f/f^ mice receiving bilateral injection of either control or synapsin-Cre.EGFP virus in the VTA show conditioning responses to exercise, as determined by conditioned place-preference. Conditioning was similar in males (right) and females (left). **B,** Control virus-injected exercised IGF-1R^f/f^ mice show significantly better mood than synapsin-Cre-injected mice, as determined by significantly higher time spent in the open arm of the elevated plus maze. Males (right) and females (left) behaved similarly (n= 8-9 per group; *p<0.05, **p<0.01 and ***p<0.001).

## Discussion

It has been shown that the liver produces hormones that regulate food intake by acting not only on the hypothalamus but also on reward circuitries (Soberg et al., 2017). The present results indicate that circulating IGF-1, produced mainly by the liver (Yakar et al., 1999), modulates physical activity acting in part through hypothalamic orexin neurons and the VTA, a component of the reward system (Morales and Margolis, 2017). Collectively, these findings demonstrate that mice deficient in serum IGF-1 exhibit reduced spontaneous activity, which can be rescued by either exogenous IGF-1 or dopaminergic drugs. Considering that: (1) IGF-1 stimulates VTA neurons, (2) IGF-1 receptors are present in VTA neurons, and (3) downregulation of IGF-1 signaling abolishes the mood-regulatory effects of exercise, we conclude that circulating IGF-1 - predominantly liver-derived - modulates VTA function in mediating the rewarding aspects of physical activity.

Additional observations in mice with blunted IGF-1R activity in orexin neurons (Firoc mice) indicate that part of IGF-1 actions for motivating physical activity are exerted by modulating orexin neurons at the lateral hypothalamus, a group of cells that project to VTA neurons (Baimel et al., 2017). Of note, administration of a dopaminergic agonist to Firoc mice did not normalize their spontaneous physical activity, suggesting that IGF-1 modulation of orexin cells is required independently of dopaminergic neurotransmission. Moreover, Firoc mice show relatively lower reduction of spontaneous physical activity compared to LID mice. Based on these two observations, we speculate that circulating IGF-1 acts on different subpopulations of VTA neurons (Lammel et al., 2014; Beier et al., 2019) through direct and indirect pathways. The indirect pathway includes IGF-1 input through orexin neurons that in turn modulate a specific subset of VTA neurons. In the absence of IGF-1-modulated orexinergic input, this VTA subpopulation is no longer affected by physical activity (Figure 7), as Firoc mice show lower activation of VTA neurons after exercise, compared to littermates. The orexin-VTA pathway involves differential dynorphin and orexin actions on VTA neurons (Baimel et al., 2017), as both neuropeptides are secreted by orexinergic neurons (Chou et al., 2001), and includes not only synapses onto DA neurons, but also onto GABAergic neurons in the VTA (Balcita-Pedicino and Sesack, 2007). A second VTA subpopulation of neurons may be directly activated by IGF-1 (Pristera et al., 2019), without receiving any orexinergic input. We speculate that this second subpopulation modulates physical activity through the IGF-1/orexin-dependent VTA population, since the dopamine agonist acts only in LID, but not in Firoc mice (Figure 7). In addition, this second VTA subpopulation may innervate distinct downstream areas (de Jong et al., 2022). Indeed, functional impact of local connections between different VTA subpopulations have been shown in relation to reward (Morales and Margolis, 2017). In agreement with these observations, we previously documented that physical activity stimulates the entrance of serum IGF-1 into the brain (Carro et al., 2000), and that circulating IGF-1 directly modulates hypothalamic orexin neurons (Zegarra-Valdivia et al., 2020).

**Figure 7:**
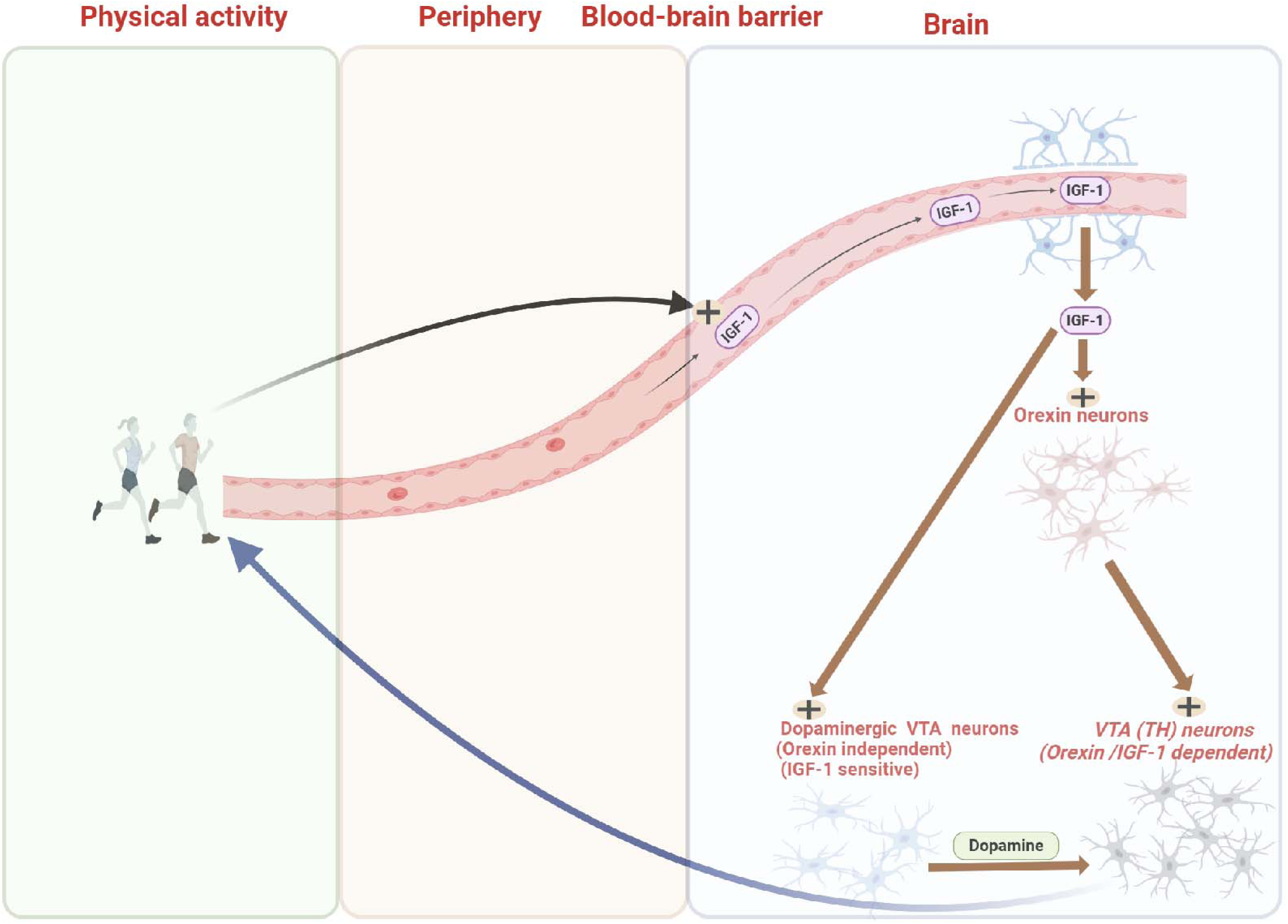
Schematic representation of the relationships between circulating IGF-1, orexin, and VTA neurons to promote physical activity. After physical activity is increased in response to internal or external stimuli, entrance of circulating IGF-1 into the brain is enhanced (Carro et al., 2000). Orexin neurons are activated by IGF-1 (Zegarra-Valdivia et al., 2020), and in turn, a subpopulation of monoaminergic (TH^+^) neurons in the VTA are stimulated by this IGF-1/orexin pathway to regulate motor output. This VTA subpopulation is not stimulated in response to exercise when IGF-1/orexin input is missing (Firoc mice). A second IGF-1-sensitive / orexin-independent subpopulation, -within at least five subpopulations of dopaminergic VTA neurons (Lammel et al., 2008), is involved in motivational motor output acting onto the first orexin-dependent subpopulation through a dopaminergic-sensitive (DR1 agonist-sensitive) pathway.

Based on these observations, we speculate that physical vigor, as reflected by muscle/bone mass, may be centrally gauged by conveying information through serum IGF-1 to brain regions involved in controlling physical activity (orexin neurons) and its emotional processing (VTA neurons). As discussed elsewhere, the rewarding effects of exercise are probably related to the key function of physical fitness in survival (Erikssen, 2001).

Proper integration of body and mind is paramount for well-being, with inner bodily sensations impinging on emotions in a continued crosstalk (Craig, 2002). An important arm of this body-to-brain signaling is provided by hormones that modulate brain function in multiple ways. The best-studied peripheral-to-brain loop is probably the one involved in energy homeostasis/fat mass, where hormones such as insulin, and many others produced by different tissues (leptin, ghrelin…etc), exert fine regulatory actions in multiple brain nuclei involved in the central regulation of energy partitioning (Kim et al., 2018). The muscle, one of the most abundant tissues of the body, should also inform brain centers to maintain physical activity according to its resources. Although muscle-derived myokines are known to influence central nervous system function (Wrann et al., 2013), and the muscle-brain connection has garnered increasing attention (Rai and Demontis, 2022) the hormonal mechanisms by which muscles and other tissues, such as bone, communicate vigor-related signals to the brain remain poorly understood. Our findings help address this knowledge gap by revealing a potential IGF-1/orexin/VTA regulatory loop that may coordinate physical activity with muscle mass in a positive feedback cycle. Specifically, we propose that exercise enhances IGF-1 signaling to orexin neurons, which then promote muscle tone, movement motivation, and metabolic balance – consistent with their established role in integrating peripheral glucose metabolism (Yamanaka et al., 2003).

A relatively recently incorporated partner in the body-brain dialog is the gut microbiome, that modulates higher brain function in different ways (Mayer et al., 2015). Interestingly, the microbiome increases serum IGF-1 levels probably through the production of short chain fatty acids (SCFAs) (Yan et al., 2016). Thus, we speculate that the microbiome also participates in modulating the information about physical vigor to the brain acting through neuroendocrine mediators (Cussotto et al., 2018), such as IGF-1. Based on the present observations, we speculate that microbe-originated metabolites such as SCFAs upregulating serum IGF-1 will induce positive mood, as recently found for depressive mood regulation by the microbiome (Chevalier et al., 2020).

These observations may provide an explanation for individual differences in willingness to exercise that could result in faulty long-term adherence to intervention programs of physical activity, a currently advocated therapeutic strategy for multiple conditions (Nunan, 2016). Since exercise behavior may be in part genetically determined (Herring et al., 2014), it could be related to the ability of IGF-1 to signal onto orexin/VTA neurons. Thus, individuals with lower IGF-1 input to these neurons would show less motivation to exercise. Appropriate serum IGF-1 input to orexin/VTA neurons will assure a proper level of physical activity, preventing in this way muscle mass loss, as seen in age-associated sarcopenia. We speculate that the latter may be in part due to inappropriate IGF-1 input to orexin neurons in old age (Zegarra-Valdivia et al., 2022).

A limitation of this study is that we used only female mice, as lower spontaneous running activity in males made it difficult to establish a significantly different behavior with the respective control males. Thus, the mechanisms described herein may be limited to females.

In summary, our results indicate that serum IGF-1, which reflects muscle and bone functioning, enters the brain in response to exercise and is involved in its rewarding effects by modulating VTA neurons through a direct and indirect pathway. The latter involves orexin neurons in the hypothalamus. Further work is needed to more accurately identify VTA circuitries affected by IGF-1 and orexin. Indeed, local IGF-1 is known to participate in modulation of VTA circuits (Pristera et al., 2019) while orexin projections to VTA have been started to be charted (Baimel et al., 2017).

## Materials and Methods

### Animals

The following mouse models were used; 1) adult female Firoc mice lacking IGF-1R in orexin neurons (25-32 g, 4-9 months old; Cajal Institute, Spain) and their littermates, obtained as described elsewhere in detail (Zegarra-Valdivia et al., 2020); 2) LID female mice (liver IGF-1 deficient, congenic with C57BL/6J mice), described before (Yakar et al., 1999); 3) IGF-1R^fl/fl^ mice (Jax Labs, B6,129 background; Jackson Labs; stock number: 012251) and 4) C57BL/6J female and male mice (Janvier Labs, France) of similar ages. Mice were housed in standard cages (48 × 26 cm²) with 4–5 animals per cage under controlled conditions: a temperature of 22°C, a 12 h/12 h light-dark cycle, and *ad libitum* access to a pellet rodent diet and water. All experimental procedures were conducted during the light phase (13:00–17:00) in compliance with European guidelines (Directive 2010/63/EU) and were approved by the relevant Bioethics Committees (Madrid Government, Proex 112/16; University of the Basque Country, M20-2022-009)

### Behavioral tests

#### Running wheel activity

We use a standard polycarbonate cage with a 25 cm diameter running wheel made of stainless steel and low friction connected to a magnetic counter that measures each turn (Cibertec, Spain). We use these cages in two sets of experiments, in short-term running (2 weeks), or in long-term running (6 months). The total amount of running is displayed as total revolutions or running speed, respectively. Running speed was determined for each bout by multiplying the number of revolutions of the wheel in that bout by the wheel circumference and dividing by the bout length. Control sedentary mice were kept in the same cages but with a locked running wheel. *Rotarod*: Animals were familiarized with the device two days before the test. On that day, 4 trials were carried out in which the mouse was placed on the rotating rod, and the speed gradually increased from 4 to 40 rpm for 5 minutes, with 10 minutes between each test. Time was recorded while mice hold on the rod until they fell or took a full turn. The performance of each mouse was obtained by making the average of the four trials (Al-Mahdawi et al., 2006). Both sexes were included.

#### Open Field

Animals were introduced in a space of 42 x 42 x 30 cm (AccuScan Instruments) for 10 minutes. Time in the center, as well as the total distance traveled, was quantified automatically by the Versamax program (AccuScan Instruments). Time in the middle of the open field was used as a measure of basal anxiety (Rodgers et al., 1997). Both sexes included.

#### Running Reward Conditioned Place Preference

The conditioned place preference (CPP) is a standardized tool used to study motivational effects of drugs and non-drugs interventions. In this protocol, we used the running wheel as a reward stimulus in mice, as described elsewhere (Fernandes and Fulton, 2016). Briefly, animals were free to run for 3 weeks in horizontal running wheels in their home cages. After that, mice were subjected to a pretest CPP task using a three-chamber maze; a white compartment with black dots, a white compartment with gray vertical lines, and separated each by a neutral center black compartment. Animals spent 5 minutes in each compartment for habituation, and then 15 min of exploring the three chambers. We measure the total time spent in each chamber and use the preference chamber as non-rewarding, and the non-preference as the rewarding chamber. The next day we started 14 days of conditioning program of access to the running wheels or not, interpolated. Mice had 2h access in their home cages to a running wheel (paired trial), or a locked running wheel (unpaired trial), and were then confined to either the black or white compartments of the CPP apparatus for 30 min. On day 15, animals were put in the apparatus for the post-test, again for 15 min of free exploring. The proportion of time spent on the paired side (running chamber) was compared to the total time obtained during the pretest (non-preference chamber).

For double orexin/c-fos or TH/c-fos immunostainings, a sub-group of animals were submitted to free running wheel for 1h for 5 days, as the previous conditioning context. The last day, 2h after running, mice were sacrificed, and their brains processed for immunohistochemistry

#### Social behavior

Social behavior encompasses both rewarding and motivational processes (Trezza et al., 2011; McCall and Singer, 2012). To assess social affiliation and novelty preference, we adapted an established three-chamber paradigm (Kaidanovich-Beilin et al., 2011). Briefly, each mouse was placed in a three-compartment arena (one central and two lateral chambers). To evaluate social affiliation (the tendency to interact with conspecifics), we introduced a stranger mouse in one lateral chamber (enclosed in a wire grid) and left the opposite chamber empty. Chamber assignments were alternated between trials to control for potential side biases. The test mouse was allowed to explore for 10 minutes, during which direct interaction time (sniffing, following, or close proximity) was recorded. Afterward, the arena was cleaned with ethanol, and the mouse was returned to the central chamber for the social novelty/preference test. The previously encountered stranger mouse (”familiar mouse”) remained in its original chamber, while a novel stranger mouse was introduced in the previously empty chamber. The test mouse was again allowed to explore freely, and interaction times with both the familiar and novel stranger were recorded.

#### Inverted screen test

This test consists of a 43 cm square of wire mesh with 12 mm squares of 1mm diameter. The screen is surrounded by a 4 cm deep wooden beading to prevent that the mice climb to the other side of the mesh. We place the mouse in the center of the wire mesh screen and rotate it for 2 min or until it falls down. The test is used to study the muscular strength of the four legs of the animal. Both sexes included. *Weight test:* This test includes seven weights (20-95 grams). Mice are hold by the base of the tail and allowed to grasp the different weights. A hold of three seconds is the criterion. If the mouse drops the weight in less than 3 sec, the time is recorded. Both sexes included.

#### Forced Swim Test

To assess depressive-like behavior, we use the forced swim test (Au - Can et al., 2012). Mice were placed in a glass cylinder (12 cm diameter, 29 cm height) containing water maintained at 23°C (15 cm depth to prevent escape). Each animal was subjected to a 6-minute forced swim test, during which behavioral responses (active swimming vs. immobility/floating) were recorded. The final 4 minutes of the session were analyzed to quantify total immobility time, defined as passive floating with only minor movements necessary to keep the head above water (Munive et al., 2019). *Elevated Plus Maze:* Anxiety-like behavior was tested in an elevated plus maze (EPM) consisting of two open arms (30 × 5 cm, no walls) and two enclosed arms (30 × 5 × 15.25 cm, surrounded by walls), arranged in a plus-shaped configuration and elevated 40 cm above the floor. Each animal was placed in the central platform (5 × 5 cm) facing an open arm and allowed to explore freely for 5 minutes. Behavior was recorded using an automated video-tracking system (Video Tracking Plus Maze Mouse; Med Associates, USA), which quantified time spent in open vs. closed arms, and number of entries into each arm. An increased preference for enclosed arms (reduced open-arm exploration) was interpreted as anxiety-like behavior, consistent with established EPM paradigms (Zegarra-Valdivia et al., 2019).

### Evaluation of muscle mass

Computerized tomography (CT) was used to measure muscle mass using procedures described in detail elsewhere (Pasetto et al., 2018). Briefly, mice were anesthetized with Isoflurane (Zoetis)5% for induction, 1.5% for maintenance via nose cone (David Kopf Instruments, France). Animals were positioned prone with limbs lateral from the torso for a uniform CT acquisition. Image acquisitions were performed using the Albira II SPECT/CT (Bruker BioSpin PCI) with 600 projections. The X-ray source was set to a current of 400 μA and voltage of 45 kVp. Images were reconstructed using the FBP (Filtered Back Projection) algorithm via the Albira Suite 5.7., re-constructor using “Standard” parameters. These combined acquisition and reconstruction settings produce a final image with 125 μm isotropic voxels. Images were segmented using PMOD v3.3 software according to tissue density-first for total volume. The distance from the upper extremity of the tibia to the medial malleolus (length of the tibia, L), and the perpendicular distance from the half-length of the tibia to the external margin of the hind limb muscle (thickness of the muscle, T) were measure. T and L measures were acquired by keeping the z axis fixed on the plane where the patella and the upper extremity of tibia were clearly identified. The Index of Muscle Mass (IMM) was defined as the ratio between T and L.

### Drug administration

Alzet osmotic mini-pumps (Model 1004; USA) were used for chronic administration of hIGF-1 (Pre-Protech, USA; 50 µg/kg/day) or the vehicle (saline). Pumps were implanted subcutaneously between the scapulae following the manufacturer’s instructions. Treatment lasted 4 weeks. In acute dosing, IGF-1 was dissolved in saline and intraperitoneally (ip) injected (1µg/g body weight). Mice were processed for immunocytochemistry (see below) two hours after ip IGF-1 injection, as described below. SKF82958 (Santa Cruz Biotechnology), a Dopamine Receptor 1 agonist, was administered for 2 weeks at 0.1mg/kg, ip, 30 minutes before the dark phase. Control mice received saline. Clozapine-N-oxide (CNO) from Tocris at 2mg/kg dissolved in saline 0.9% was injected intraperitoneally 40 min before test sessions.

### Immunocytochemistry

Animals were anesthetized with pentobarbital (50 mg/kg) and perfused transcardially with saline 0.9% and then 4% paraformaldehyde in 0.1 M phosphate buffer, pH 7.4 (PB). Coronal 50-μm-thick brain sections were cut in a vibratome and collected in PB 0.1 N. Sections were incubated in permeabilization solution (PB 0.1N, Triton X-100, NHS 10%), followed by 48 hours incubation at 4°C with primary antibody (1:500) in blocking solution (PB 0.1N, Triton X-100, NHS 10%). Antibodies used in this study include rabbit polyclonal c-Fos (Abcam ab190289), orexin polyclonal goat antibody (Santa Cruz 8070), orexin polyclonal rabbit antibody (Abcam ab 6214), anti-NeuN (Invitrogen MA533103), monoclonal G5 anti-IGF-1 receptor α (ref. sc-271,606, Santa Cruz), and monoclonal tyrosine hydroxylase (TH) mouse antibody (Millipore, MAB318). Followed by goat polyclonal - Alexa Fluor (488 / 594) as secondary antibody (1:1000). Finally, a 1:1000 dilution in PB of Hoechst 33342 was added for 5 minutes. Slices were rinsed several times in PB, mounted with gerbatol mounting medium, and allowed to dry. Omission of primary antibody was used as control.

### RNAScope

For *in situ* hybridization, brains were fixed for 2 hours at room temperature (RT), and then cryoprotected in 30% sucrose and embedded in octanol (OCT) to store at -80°C until sectioning in the cryostat (CM1950, Leica Microsystems) at 10-15 μm thick. The sections were collected directly from super-frost ultra plus slides (Thermo Scientific, #10417002) and stored at -80°C until processing using RNAScope (2.5 HD Detection kit—Red; #322350; ACD, USA) with an IGF-1R exon 3-specific probe (Mm-Igf1r-O1, #577471) combined with immunocytochemistry with anti-NeuN antibody to confirm the deletion of exon 3 in VTA cells after virus injection, as described before (Fernández de Sevilla et al., 2022).

### Cell Image Analysis

Images were taken from the VTA and LH in a confocal microscope (Leica, Germany) using 20X magnification for c-fos. For double-stained TH/c-fos or orexin/c-fos cell counting, four sections per animal were scored using the MATLAB analysis (Dima et al., 2011) by a blinded experimenter. First, confocal images were segmented via k-means clustering with the number of clusters set to 5 (k = 5) using MATLAB to obtain binary masks for each channel dividing background from the foreground. Red channel masks (TH or Orexin antibody) were further processed by individualizing unconnected regions, putative neurons, and eliminating isolated components with areas smaller than 25 μm^2^. The number of neurons in each mask was manually assessed. Green channel masks (c-fos antibody) were also visually inspected to ensure proper segmentation. Red and green channels were superposed to study the area of co-localization. For each segmented neuron (red channel), the fraction of its area that co-localized with c-fos antibody (green channel) was measured as an indirect quantification of the amount of c-fos present in that neuron. We referred to this fraction as the co-localization index. The co-localization index of each image was computed as the mean co-localization index of the neurons present in that image. Lastly, the final co-localization index for each condition was computed as the mean co-localization index of all the images belonging to that condition. Images with only secondary antibody staining were used to check specificity of the secondary antibody (Burry, 2011), and segmented as described above. Specificity was evaluated by measuring the segmented and the number of segmented neurons, comparing the results with tissue images that underwent a complete immunohistochemistry protocol.

### Viral constructs

For chemogenetic experiments using DREADD, a viral construct (pAAV-hSyn-DIO-hM4D(Gi)-mCherry ; AAV5; 8.6 x 10^12^ viral infective units/ml) was locally injected bilaterally (AP= -1.4; ML= ± 0.9; DV= -5.4) to target orexin neurons in Orexin-Cre mice. As a control virus, we used pAAV-hSyn-DIO-mCherry (AAV5). Both viral constructions were obtained from Addgene. Clozapine N-Oxide (CNO, 2mg/kg dissolved in saline 0.9%) was administered ip and 40 min later behavior was assessed, CNO efficacy in orexin neurons was previously confirmed in acute slices obtained from virus-injected mice (Fernandez de Sevilla et al., 2020). Synapsin-driven Cre, AAVPHP.B-SYN-CRE and AAVPHP.B-CMV-EGFP (as control) viral constructs were bilaterally injected into the VTA (AP: -3.5, L:0.35, DV:4.5) in control mice. After four weeks, animals were used.

### Surgery

For surgeries, mice were anaesthetized with isoflurane as above, and placed on a stereotaxic frame (Stoelting Co) on a heating pad and tape in their eyes to protect them from light. For viral expression, mice were injected with a 5 µl Hamilton syringe bilaterally into the orexin nuclei (AP= -1.4; ML= ± 0.9; DV= -5.4) or VTA (AP: -3.5, L:0.35, DV:4.5) 4 weeks before experiments. Three hundred nl were bilaterally infused at a rate of 100 nl/min and the syringe withdrawn 10 min later.

### Statistics

Statistical analysis was performed using Graph Pad Prism 6 software (San Diego, CA, USA). Depending on the number of independent variables, normally distributed data (Kolmogorov-Smirnov normality test), and the experimental groups compared, we used either Student’s t-test or two-way ANOVAs by Sidak’s multiple comparison test. For non-normally-distributed data, we used the Mann-Whitney U test to compare two groups, Kruskall-Wallis or Friedman test, with Dunn’s multiple comparisons for multiple groups. The sample size for each experiment was chosen based on previous experience and aimed to detect at least a p<0.05 in the different tests applied, considering a reduced use of animals. Results are shown as mean ± standard error (SEM) and p values coded as follows: *p< 0.05, **p< 0.01, ***p< 0.001.

## ACKNOWLEDGMENTS

We are thankful to M. García for technical support.

## AUTHOR CONTRIBUTIONS

J.A. Zegarra-Valdivia designed and conducted experiments, prepared figures, results, and wrote part of the manuscript, J. Fernandez, K. Suda, M.Z. Khan, and M.E. Fernandez de Sevilla conducted experiments, J. Pignatelli, J. Esparza, and S. Diaz helped with experiments. I Torres Aleman designed the study and wrote the manuscript.

