## Supplementary Figure Legends for "INTEROCEPTIVE INFORMATION OF PHYSICAL VIGOR THROUGH CIRCULATING INSULIN-LIKE GROWTH FACTOR 1"

**Supplementary Figures**

**Supplementary Figure 1. A,** Spontaneous physical activity in male WT mice is lower than in females (males=9, females=15; ***p<0.0001, Two-Way ANOVA). **B,** Total running differences between males and females (males=9, females=15; ***p<0.0001, Mann-Whitney U test). **C,** LID mice selectively reduced running behavior in the active phase (n=5 per group; ***p<0.0001, Two-Way ANOVA). **D,** Number of double-labelled c-fos/orexin^+^ neurons in sedentary (n=24) and exercised control mice (p<0.01, n=22). E, Representative pictures of double-labeled c-fos/orexin neurons in WT mice under sedentary and exercise conditions. Bars are 50 µm. **F,** Number of orexin neurons in control sedentary (n=24) and exercised mice (n=22).

**Supplementary Figure 2. A-B,** Number of orexin^+^ neurons in LH did not differ significantly between experimental groups: littermates=2189 ± 511.6 vs Firoc=1668 ±482.2; p=0.4359, (Mann-Whitney test), and controls=553.2 ± 90.91 vs LID=641.3 ± 135.1 cells, p=0.4710 (One-Way ANOVA). **C,** No differences in body weight were seen between Firoc mice and littermates (3-4 months old, WT =18, Firoc =12, p= 0.2974, Unpaired t-test). **D,** Firoc mice and littermates have similar muscle mass, as determined by computerized tomography (CT) imaging. A representative CT scan showing the calculation of the length and diameter of the leg muscle. **E,** Quantification histograms of muscle mass index (n=5 per group; p=0.8357, Unpaired t-test). **F,** Performance in the inverted screen test, designed to measure strength in the muscles of the four legs, was similar in Firoc mice and littermates (WT=12, Firoc=18; p>0.999, Mann-Whitney Test). **G,** An additional measure of muscle strength, the weight-lifting test, also indicates similar strength in both experimental groups (WT=12, Firoc=18; p=0.8743, Unpaired t-test). **H,** Motor coordination in the rotarod test was unperturbed in Firoc mice (WT=11, Firoc=16; p=0.6359, Mann-Whitney Test). **I,** Deambulation in the open field test was similar in Firoc and littermates (n=10 per group; p=0.8357, Mann-Whitney Test). We use female and male mice in these behaviors, no differences were found between sexes.

**Supplementary Figure 3. A,** Timeline of behavioral experiments. Behavioral analyses started with a social interaction test (affiliation and novelty/preference). Next, a new set of animals were submitted to running for 3 weeks. After that, a pre-CPP test was performed. Then, animals were conditioned to the running wheel for 14 days. On day 15 a post-CPP was performed. Finally, the EPM was carried out. A period of 4 days was left between behavioral tests. **B,** Quantification histograms of double-labeled orexin/TH^+^ neurons in WT mice under sedentary and exercise (free-wheel running) conditions show significant differences (n=10 fields in the LH area, 3 mice per group; *p=0.0137, Mann Whitney test). **C,** Representative pictures of WT mice submitted to voluntary wheel running or sedentary conditions. Sedentary mice show a reduced number of double-labeled orexin (red)/TH (green) cells in the VTA. Bars are 50 µm.

**D,** Number of TH^+^ neurons in VTA did not differ between experimental groups (WT sedentary=686 ± 224.7, exercise=746.2 ± 144.4; p=0.8245, Unpaired t-test test). **E,** Time spent in the open arms of the EPM in sedentary (n=15) and exercised control mice (n=25); *p<0.05).

**Supplementary Figure 4. A,** Number of TH^+^ cells in VTA did not differ between saline (n=1170 ± 245.8) and IGF-1-injected mice (n=775.7 ± 193.4), p=0.2472, Unpaired t-test. **B,** Number of TH^+^ cells in VTA did not differ between Firoc and wild type (WT) littermates (WT=2189 ± 511.6, Firoc=1668 ±482.2; p=0.4359, Mann-Whitney test). **C,** Number of TH^+^ cells in VTA are similar between LID and littermates (WT=1193 ± 94.93, LID=1257 ± 104.3; p=0.6571, Mann-Whitney test). **D,** Administration of the D1 agonist SKF82958 (0.1mg/kg, ip) to LID mice significantly increased their spontaneous activity compared to saline-injected controls and naïve LID mice (LID saline=10; LID D1 agonist=9; LID naïve= 8; ***p<0.001). **E,** Total running along treatment time was increased in SKF82958 -treated LID mice, compared to naïve (**p=0.0052) and LID saline (**p=0.0042), Kruskal-Wallis test). Daily injection of saline reduced spontaneous activity as compared to naïve LID mice probably because the animals were stressed by the procedure.

**Supplementary Figure 5. A,** IGF-1 receptors are expressed by TH^+^ neurons in the VTA. Representative photomicrograph showing double TH^+^/IGF-1R^+^ cells in the mouse VTA. Note the punctuated pattern of the IGF-1R, as documented before using the same antibody ^67^. Bar is 100 µm. **B,** TH^+^ neurons (green) in the VTA immunostained with GFP (red dots) after AAV-synapsin-Cre.EGFP injection. Bar is 50 µm. **C,** Truncation of IGF-1R was confirmed by in situ hybridization in brain slices containing the VTA region using an exon 3 IGF-1R probe that is lost after Cre recombination. Non-truncated IGF-1R (with the presence of exon 3 using a specific RNAscope probe, in red) was significantly decreased in VTA neurons (NeuN^+^ in green) after AAV-synapsin-Cre.EGFP injection. Histograms: double labelled NeuN/exon-3 probe cells estimated as percentage of total NeuN cells. Probe labelling was observed in neurons (NeuN+) of FIR control, but not in FIR GFAP Cre. Bars are 25 µm and 5 µm (insets). (n = 100 cells/group, Chi-Square; ***p<0.001).
