## Supplementary Figures for "INTEROCEPTIVE INFORMATION OF PHYSICAL VIGOR THROUGH CIRCULATING INSULIN-LIKE GROWTH FACTOR 1"

### Slide 1
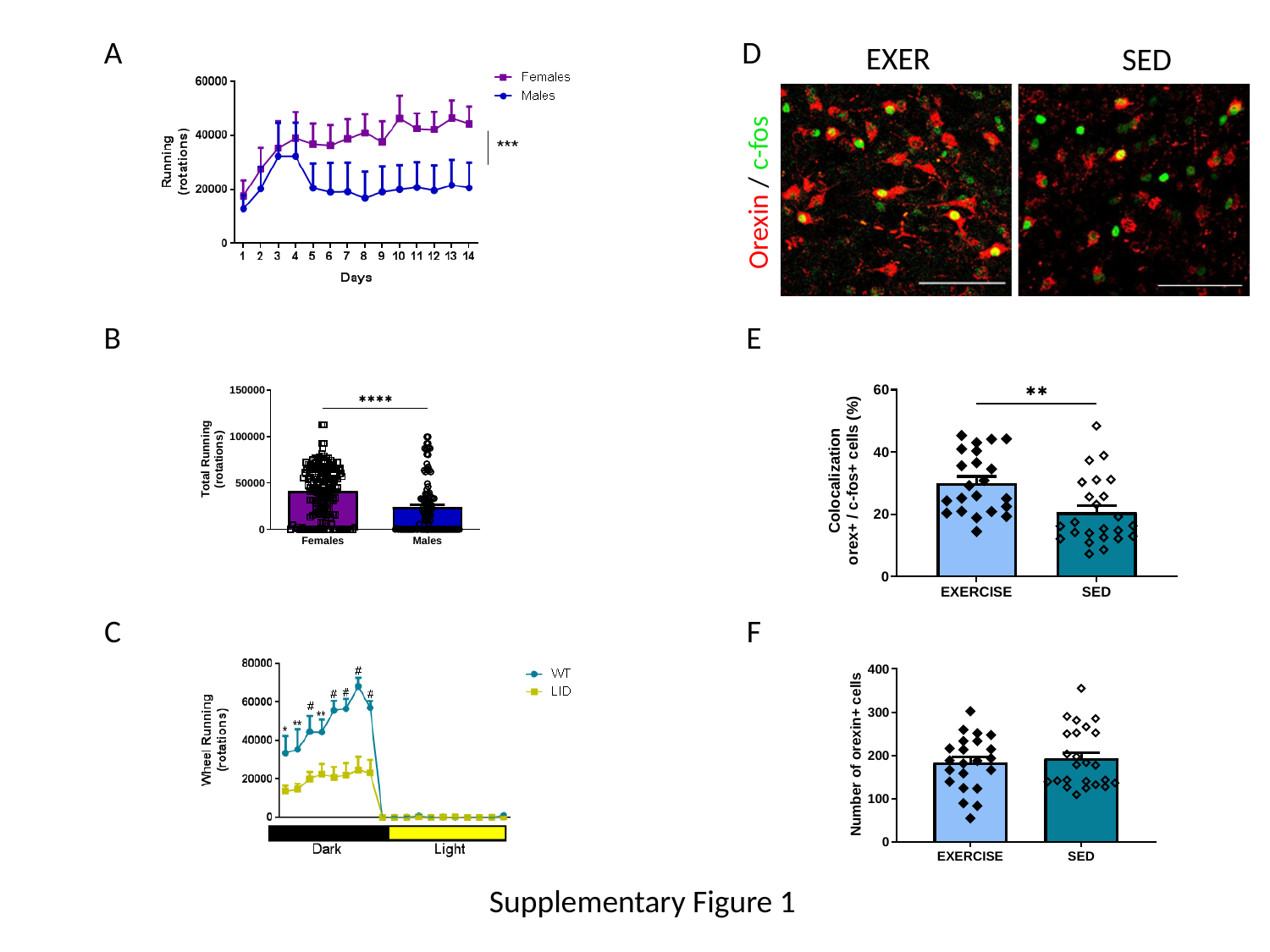

A
D
EXER
SED
Orexin / c-fos
B
E
C
F
Supplementary Figure 1

### Slide 2
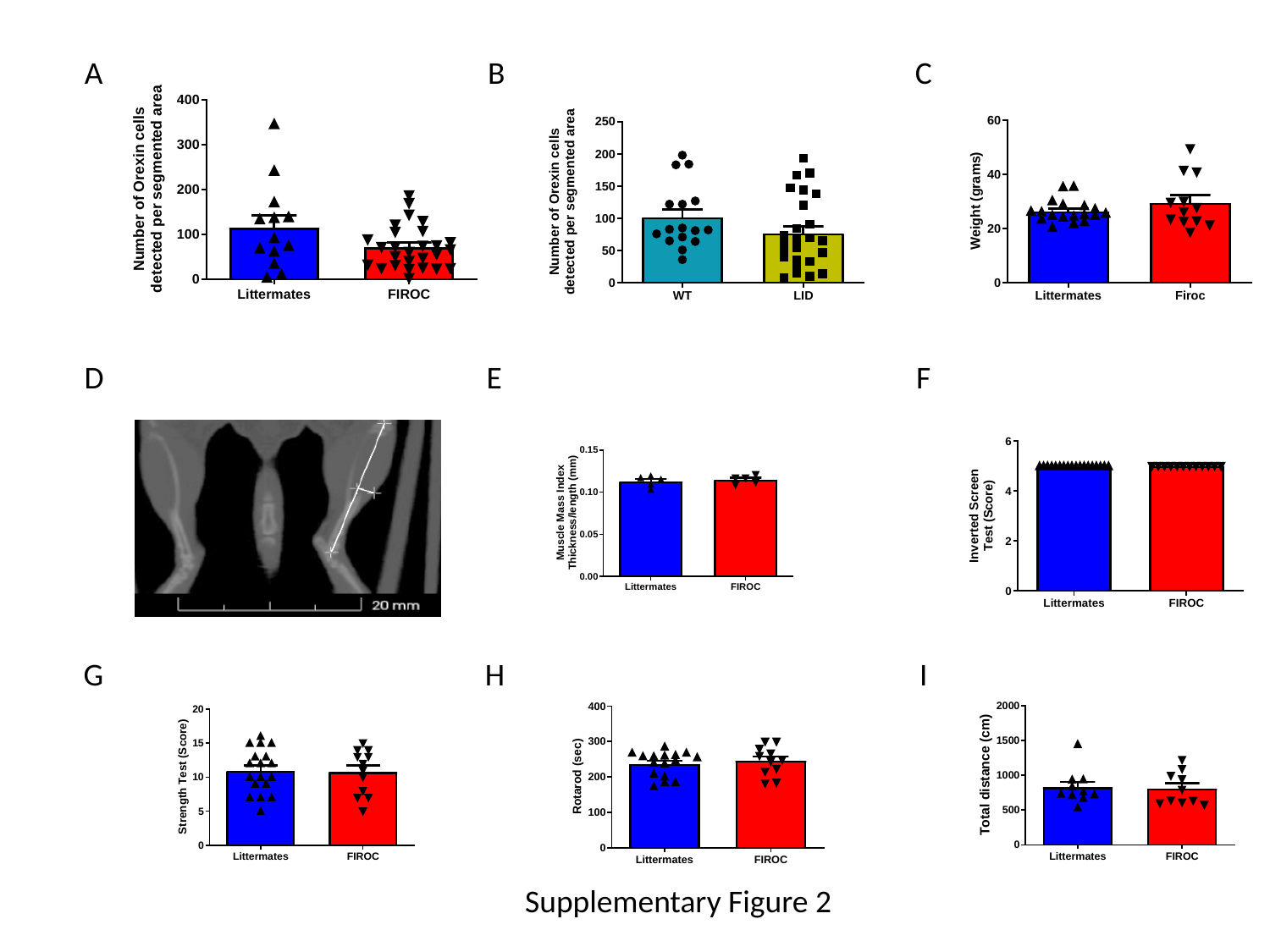

A
B
C
D
E
F
G
H
I
Supplementary Figure 2

### Slide 3
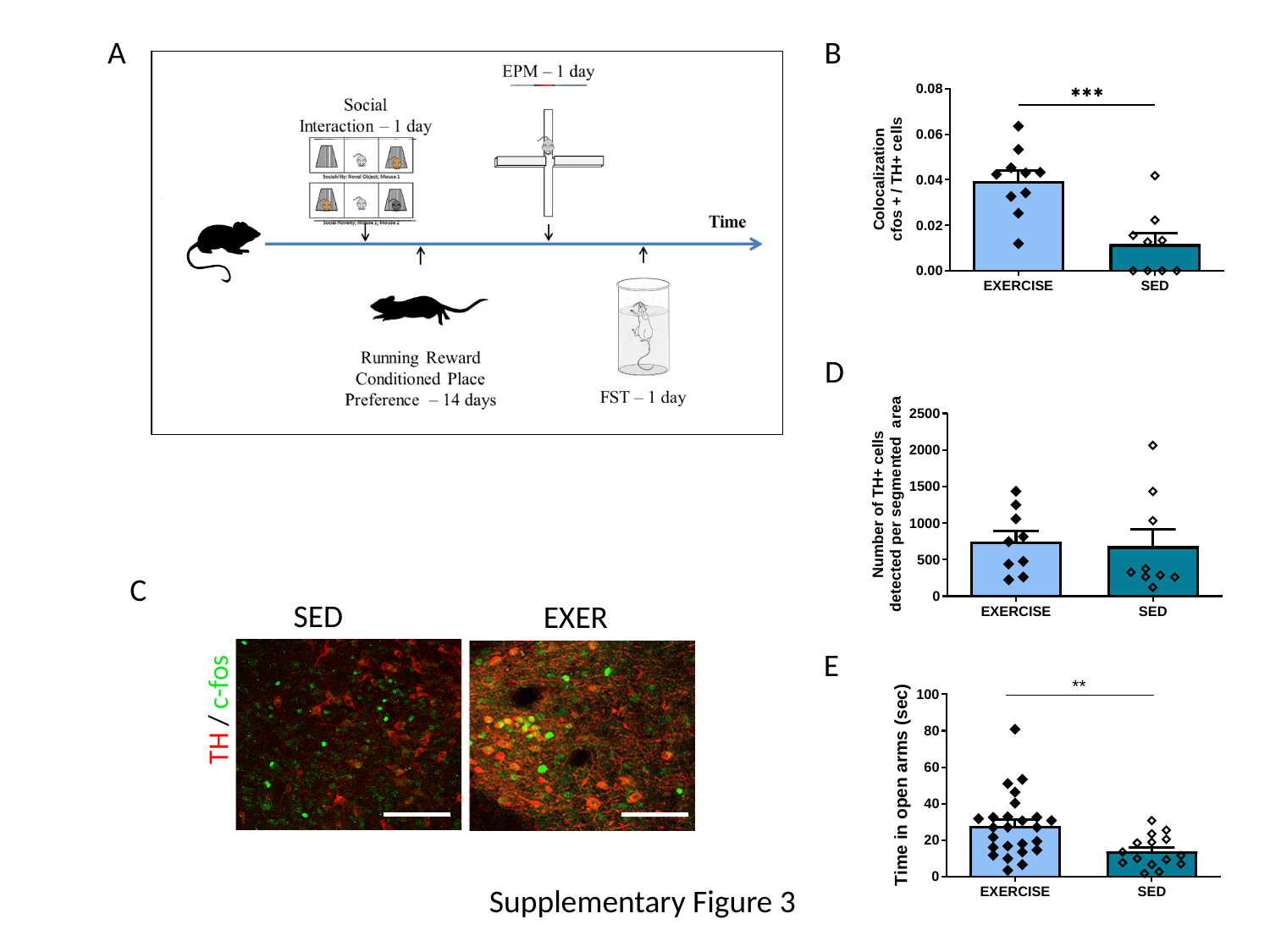

A
B
D
C
SED
EXER
TH / c-fos
E
Supplementary Figure 3

### Slide 4
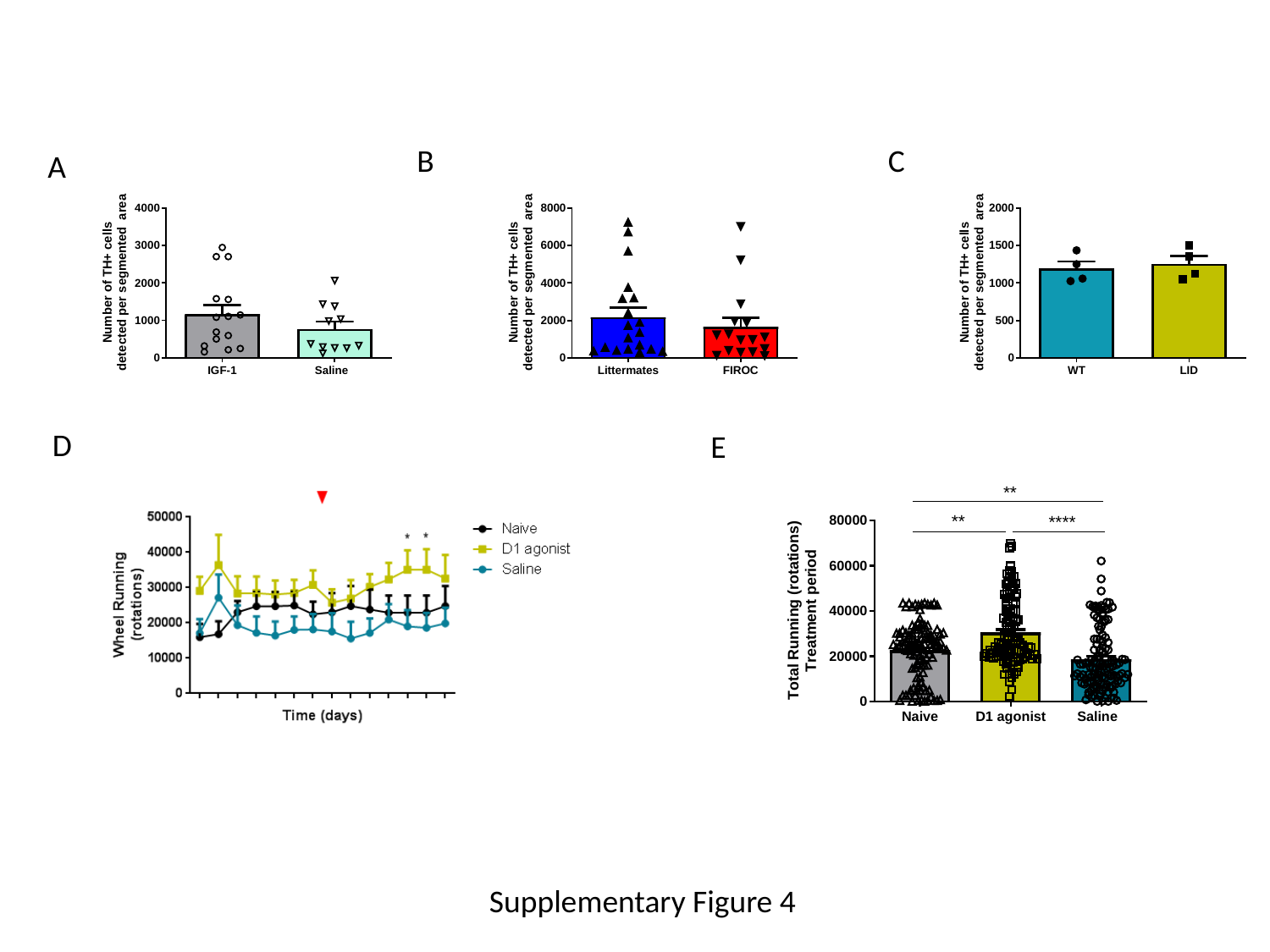

B
C
A
D
E
Supplementary Figure 4

### Slide 5
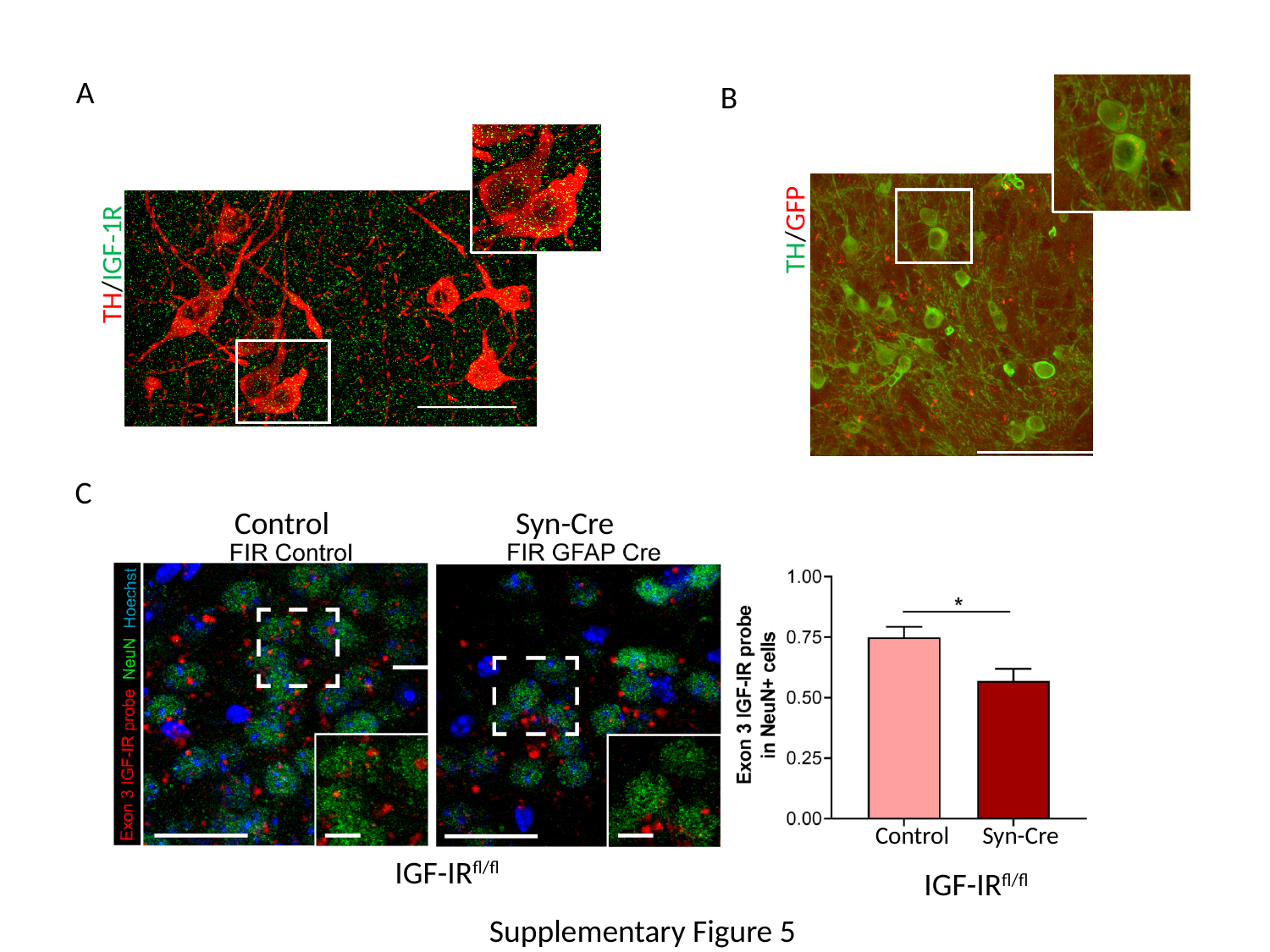

A
B
TH/GFP
TH/IGF-1R
C
Control Syn-Cre
Co
Control Syn-Cre
IGF-IR^fl/fl^
IGF-IR^fl/fl^
Supplementary Figure 5
